# A Darwinian Uncertainty Principle

**DOI:** 10.1101/506535

**Authors:** Olivier Gascuel, Mike Steel

**Author notes:** Both authors contributed equally to this work. **Correspondence to**:, Olivier Gascuel, https://research.pasteur.fr/en/member/olivier-gascuel/olivier.

## Abstract

Reconstructing ancestral characters and traits along a phylogenetic tree is central to evolutionary biology. It is the key to understanding morphology changes among species, inferring ancestral biochemical properties of life, and recovering migration routes in phylogeography. The goal is twofold: to reconstruct the character state at the tree root (e.g. the region of origin of some species), and to understand the process of state changes along the tree (e.g. species flow between countries). Although each goal can be achieved with high accuracy individually, we use mathematics and simulations to demonstrate that it is generally impossible to accurately estimate both the root state and the rates of state changes along the tree branches from the observed data at the tips of the tree. This inherent ‘Darwinian uncertainty principle’ concerning the simultaneous estimation of ‘pattern’ and ‘process’ governs ancestral reconstructions in biology. Increasing the number of tips improves the joint estimation accuracy for certain tree shapes that arise in evolutionary models, however, for other trees shapes it does not.

---

Reconstruction of the past is central to evolutionary biology (1–3). A first step is often phylogenetic reconstruction, which is central to understanding the origin, evolution and classification of species, protein families, and pathogens such as HIV, as well as for reconstructing the evolution of communities and ecosystems. However, phylogeny is not an end in itself; it is generally the support for more complete studies. In particular, one frequently reconstructs the evolution along a phylogenetic tree of ancestral characters of diverse nature, for example: molecular (4–5), phenotypic (6–7), geographical (8–12) or ecological (4–6), and these reconstructions involve differing time scales, ranging from a few years for fast evolving viruses (e.g. Ebola (12)), to dozens of millions years for higher eukaryotes (e.g. plants (4–7)). The problem has two facets (Fig. 1), which are generally combined: one may want to infer the ‘pattern’, that is, the ancestral states associated with phylogeny root and nodes, for example, the origin and migration routes of a species (9, 11) or an epidemic (10, 12); or one may aim to understanding the ‘process’ driving the character evolution and state changes, such as some adaptive mechanism or trace of positive selection (13–14). Many methods have been proposed to reconstruct the pattern. Today, one most often uses probabilistic methods based on Markovian evolutionary models with numerical parameters to be estimated from the data (1–3). These models and their parameters are mathematical representations of the evolutionary processes. We show here using information theory, mathematics and simulations, that evolutionary patterns and processes cannot generally be simultaneously reconstructed with high accuracy from extant data. This result applies even to the simplest models, and to characters commonly used in a number of current studies, to describe a wide range of evolutionary phenomena, from molecular to ecological levels. This inherent ‘Darwinian uncertainty principle’ governs ancestral reconstructions in biology.

**Figure 1.**
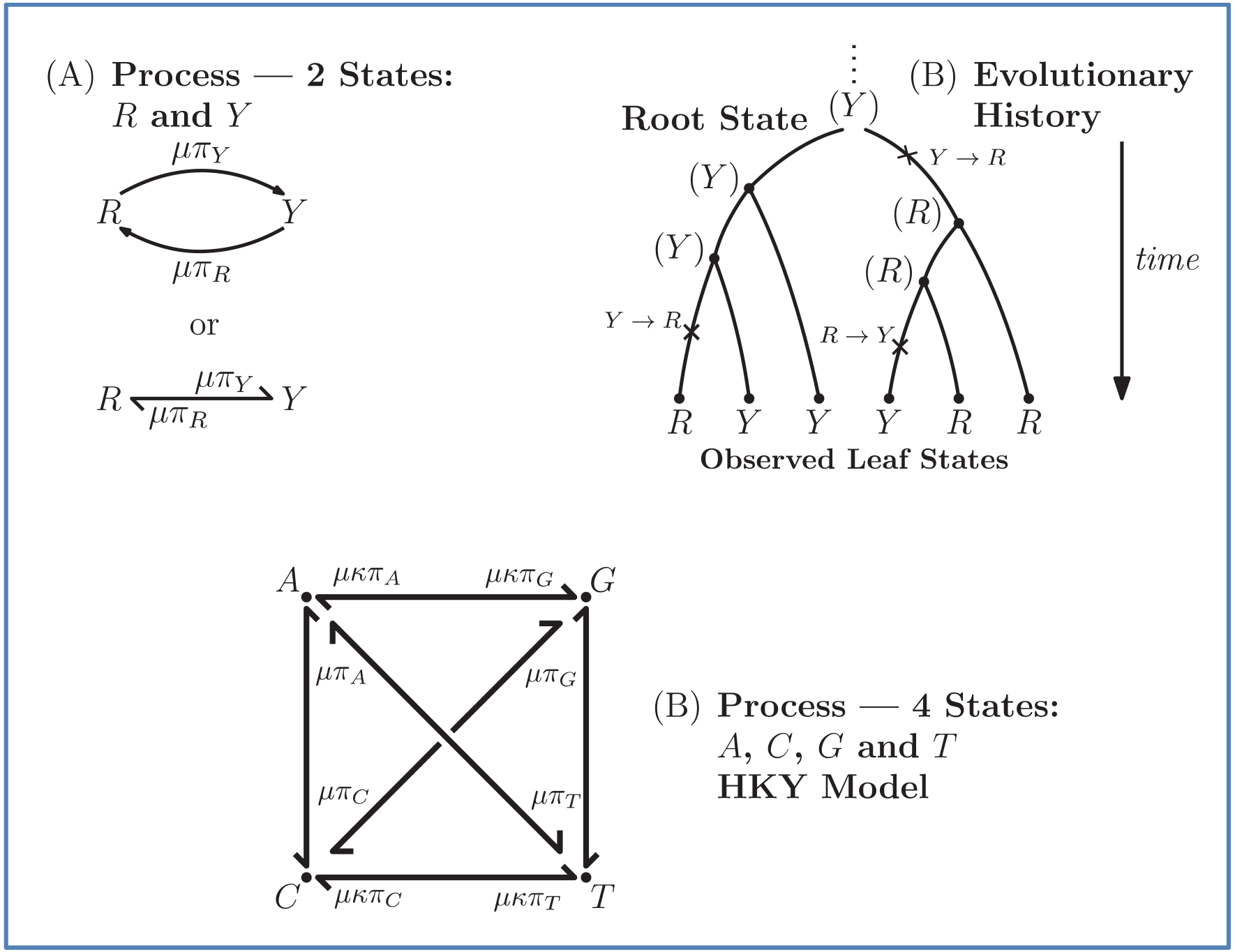
Evolutionary process and history: (A) Simple 2-state Markovian evolutionary process, where *R* swaps into *Y* with rate *μπy* and vice versa. This process has three (non-independent) parameters: the state equilibrium frequencies (*π_R_* and *π_y_*) and the global rate of evolution (*μ*). (B) This simple process acts along a phylogenetic tree, starting from the tree root with state *Y* and evolving the character state until the tree leaves, where various state values are observed. This observation (along with the phylogenetic tree) is all that is known. The goal is to estimate both the evolutionary pattern, notably the root state, and the parameters of the process. (C) More complex 4-state process, where the rate of change depends on the origin state, and not-only on the destination state. In addition to the four equilibrium frequencies attached to the four nucleotides (two purines: A and G, and two pyrimidines: C and T), the HKY model includes the parameter k, which is equal to the transition/transversion ratio. A transversion is a change from one purine to one pyrimidine and vice versa (e.g. A ↔ C). A transition does not change the nucleotide category (e.g. A ↔ G). Transitions tend to be faster than transversions (i.e. *κ* > 1). *κ* = 1 corresponds to the F81 or ‘equal-input’ model.

The Markovian evolutionary models used to reconstruct character evolution can be very simple, typically symmetrical with very few states (Fig. 1A), but the current trend is to rely on ever more complex models which can be non-symmetrical, with dozens of states (12) (and therefore hundreds of parameters), latent variables (6) and, for some models, evolution over time (7–8, 11–12). The estimation of these models is based on maximum likelihood and Bayesian approaches, and the task is complicated by the fact that there is only one realization of the process, corresponding to the observed values of the states observed at the leaves of the tree. Estimations are much simpler with DNA or protein sequences, where we assume the same model for all the characters and thus benefit from multiple information sources. To learn the most complex of these (single-character) models, one can rely on user-supplied ‘factors’, such as the degree of connectivity between two countries in phylogeography (10, 12). To predict ancestral states, one generally uses marginal, joint or posterior likelihoods of the tree node states (1–3, 15). These two components (model estimation and ancestral reconstruction) are most often simultaneous and interdependent because neither the model nor the ancestral states are generally known. Only the tree and branch lengths can be considered as known; in practice, they are usually estimated from DNA or protein sequences via a probabilistic approach, possibly including a molecular clock model to date the tree nodes and root age (2).

Theoretical work has shown the difficulty of reconstructing ancestral states even when the evolutionary model describing the state changes is fully known (16–18). If the rate of changes is too fast, the information provided by tree leaves is low and it is impossible to reconstruct the root state accurately, regardless of the estimation method. To our knowledge, there is no theoretical work on the joint estimation of evolutionary model parameters and the ancestral states. Moreover, very few simulations have been performed to verify that the complex models used in recent studies (see above) could be estimated with high reliability. We show here that it is usually not possible to accurately reconstruct both the root character state and estimate the parameters of the evolutionary model. Intuitively, if the global rate of change is low, the reconstruction of the root is easy because the root state is largely preserved along the tree branches all the way to the leaves, but then they are too few state changes to accurately estimate the relative rates of changes from one state to another; conversely, with a rapid evolution, estimating the relative rates is easier, but one cannot reconstruct the tree root.

## Mathematical Results

We first establish this finding by using mathematical results based on standard Markovian evolutionary models. For our first theorem, we consider a Markov process such as the Felsenstein 1981 model (2) for DNA (F81, also known as the Tajima—Nei model) or any other ‘equal input model’ (19) on any number of states (Fig. 1A and 1C, assuming *κ*= 1). In this model the rate of changes from state *i* to state *j* is proportional to the model equilibrium frequency of *j*, and does not depend on *i*. This model is very simple but includes the state equilibrium frequencies (to be estimated from the data), like most if not all evolutionary models used nowadays. Let *X_L_* be the observed states at the leaves (‘the data’) of a given phylogenetic tree *T* (with known branch lengths), and let *n* be the number of tree leaves. Information theory provides a precise way to formalize our first result. Let *I_ρ_* denote the mutual information between *X_L_* and the ancestral state at the root of tree *T*, and let *I_π_* denote the mutual information between *X_L_* and the state equilibrium frequency vector (*π*) of the model. The root state is sampled from π, as commonly assumed in phylogenetics. Both *I_ρ_* and *I_π_* are functions of the global evolutionary rate *μ* of character evolution.

Our first theorem (described in Methods) demonstrates that the information provided by the data obtained at the tips of an evolutionary tree concerning the ancestral root state and concerning the relative rates behave in opposite ways as a function of the global evolutionary rate *μ* More precisely, Theorem 1 says that for any tree, as *μ* increases, *I_ρ_* and *I_π_* always have consistent but opposite trends. In particular, the optimal substitution rate for estimating the ancestral root state is the worst for estimating *μ*, whereas the optimal substitution rate for estimating *π* is the worst for estimating the ancestral root state. This immediately implies a fundamental uncertainty limit on the accuracy of simultaneous estimation of both these variables.

Our second theorem (described in Methods) positively moderates this phylogenetic uncertainty principle with Yule trees (20–23), which roughly describe the shape of speciation trees. Theorem 2 shows that for Yule trees, increasing the number of tips reduces the uncertainty of simultaneous estimation. This result holds for a wide variety of evolutionary models (we can allow *any* stationary, reversible, continuous-time Markov process involving any number of states for which the rate matrix *R* has strictly positive off-diagonal entries). However, for some other tree shapes (e.g. coalescent trees (24), commonly used in population genetics, and star trees, corresponding to extreme radiations), uncertainty remains even if the number of tips tends to infinity.

## Simulation Results

To explore the behavior of evolutionary models that are more complex and realistic than F81 and equal input models of Theorem 1, we use computer simulations. The goals are to: quantify the uncertainty with both Yule and coalescent trees; measure the gain brought by a large number *n* of tree leaves and observed states; and study the accuracy of estimations with model parameters that are different from the simple equilibrium frequencies defining F81 and related models. We use the HKY model (2, 25) of DNA evolution (Fig. 1C). In addition to the four equilibrium frequencies attached to the four nucleotides, this model includes the parameter *κ*, which is equal to the transition/transversion ratio. A transversion is a change from one purine to one pyrimidine and vice versa; a transition does not change the nucleotide category. Transversions from state *i* to state *j* occur at a rate proportional to *μπ_j_*, whereas transitions occur at a rate proportional to *μκπ_j_*. Thus, the rate of changes does not only depend on the destination state, but also on the origin state, unless *κ*= 1, which corresponds to F81. HKY represents a larger and more realistic class of models, and is commonly considered as a good model of DNA evolution. While the state equilibrium frequencies can be approximated by counting the number of state occurrences on the tree leaves, *κ* is not directly observable and its estimation is expected to be more difficult than the estimation of *π*. In our simulations, a unique character was evolved using an HKY model along Yule and coalescent trees of 100 and 1,000 tips, with various values of *μ* rate, from 1/16 (very slow) to 16 (very fast). All estimations were performed using the maximum likelihood (ML) principle, which is known to be optimal (26). Additional details are given in Methods.

Though the HKY model is more general than equal input models, the results (Fig. 2) are in accordance with the uncertainty principle of Theorem 1, for both Yule and coalescent trees. With a low *μ* rate, the root state is easy to predict but estimation of the model parameters is very poor. With a high *μ* rate, predicting the root state becomes impossible, but the equilibrium frequencies (π) are well estimated. For the rate parameters (*κ* and *μ*), their estimation first improves when *μ* increases, and then becomes poorer with Yule trees and large *μ* values, due to large numbers of changes in pending branches. This makes the tip states nearly independent one from the other, which is an advantage to estimate the equilibrium frequencies, but not the rate parameters. This finding reinforces again the uncertainty principle, as with large *μ* neither the root state nor the rate parameters can be accurately estimated. With coalescent trees (having short pending branches), the estimation of *κ* and *μ* is still improving with *μ* = 16 (Fig. 2), but drops with extreme *μ* values (results not shown). As expected from Theorem 2, the accuracy of all estimations improves with Yule trees when *n* = 1,000, compared with *n* = 100. With *n* = 1,000 we observe a narrow region around *μ* = 1 (corresponding to 1 expected mutation along any root-to-tip path), where the simultaneous estimation of all parameters (including the root state) is reasonably accurate (error < 25%). However, outside this region some of the parameters are still poorly estimated. With coalescent trees, we do not observe such a region, and (as expected) the estimation of the root state has similar accuracy with *n* = 100 and *n* = 1,000. Lastly, a surprising and positive finding is that the accuracy of root state estimation is not affected by the poor estimation of the model parameters: the results are nearly the same when using the estimated parameter values (RootMLfull) and their true values (RootMLtrue), and this finding still holds with low *μ* rate when the model parameters are very poorly estimated.

**Figure 2.**
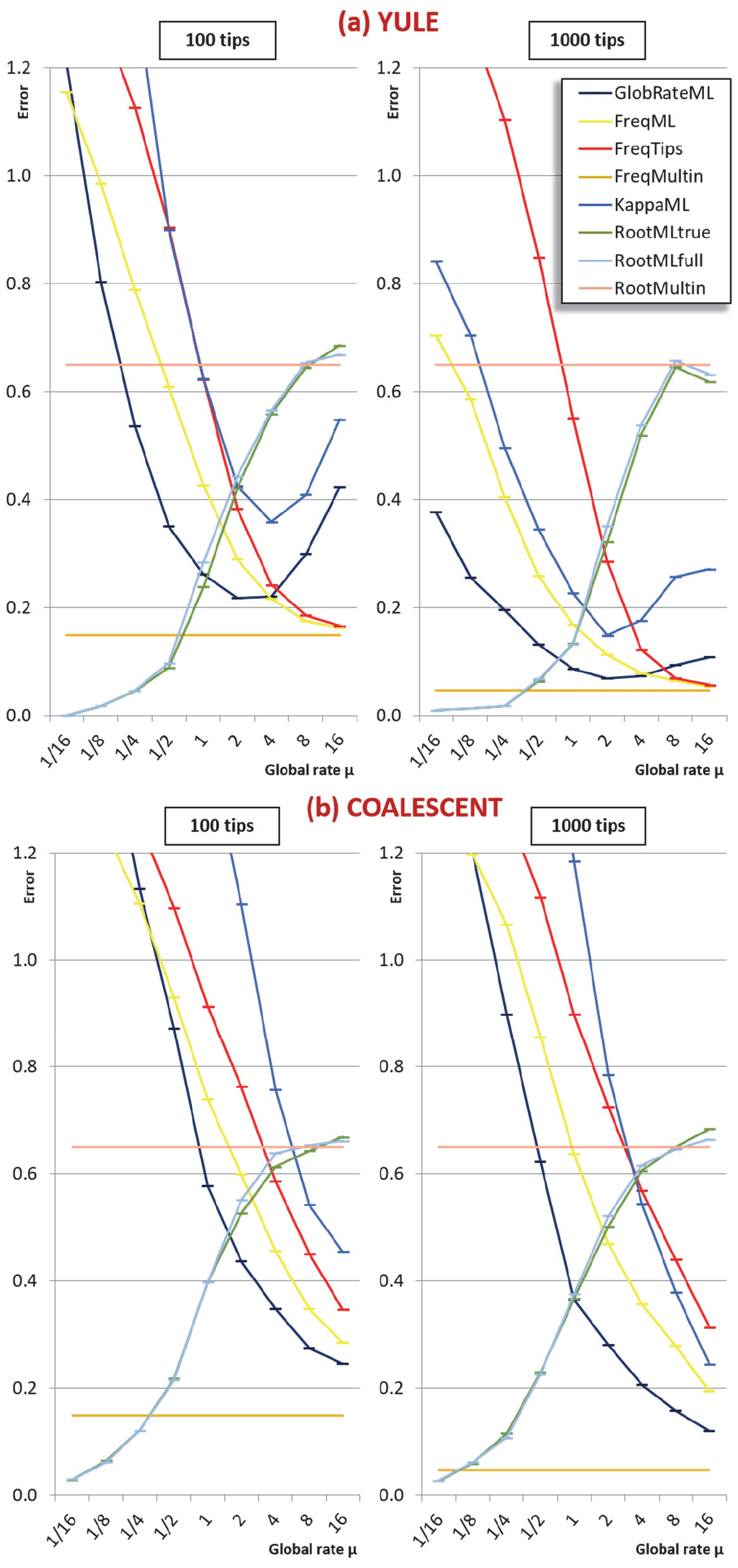
Simulation results with Yule (a) and coalescent (b) trees: Horizontal axis: value of the global rate *μ* used to simulate the data. Vertical axis: error measurements (probability of error for root state predictions; relative absolute error for other estimations; see Methods). GlobalRateML: ML estimation of the global rate μ; FreqML: ML estimation of the equilibrium frequencies; FreqTips: quick estimation of the equilibrium frequencies by counting the number of state occurrences on the tree leaves (clearly worse than ML estimation); FreqMultin: best possible estimation of the state frequencies with *n* samples, as obtained with a multinomial; KappaML: ML-based estimation of *κ* (the transition to transversion ratio); RootMLtrue: root state prediction by ML, with the knowledge of the evolutionary model used to simulate the data; RootMLfull: root stat prediction when all model parameters (*μ, π, κ*) are estimated from the data; RootMultin: root state ‘prediction’ with uninformative data, as obtained with a multinomial model (similar to a tree with very long edges).

## Discussion

Although our results (theorems, simulations) demonstrating and quantifying the uncertainty principle are obtained in simple settings, it is highly likely that with more complex models and real biological data the situation is even worse. Theorem 1 and the simulation results have a similar flavor to a fundamental principle in quantum physics – Heisenberg’s uncertainty principle - which provides an absolute lower bound on the precision of simultaneously estimating both the position and the momentum of a particle (27). Here, we take the phylogenetic analogue of ‘position’ as ‘ancestral state’, and thus ‘momentum’ (closely related to velocity) corresponds to the rates at which ancestral states change into different alternative states. In physics, increasing the mass of a particle reduces the uncertainty of jointly specifying its position and momentum; in our setting, the analogue of mass is *n*, the number of leaves. Theorem 2 shows that for certain tree shapes (Yule trees) increasing *n* also reduces the uncertainty of joint estimation. Though the models and mathematics are radically different, our results thus have a similar spirit: it is not possible to accurately estimate both the ancestry and the rate of state changes in characters commonly used in a number of current studies, to describe a wide range of evolutionary phenomena, from molecular to ecological levels.

## Methods

### Statement and sketch proof of Theorem 1

Let *T* be a rooted phylogenetic tree (not necessarily binary), for which each edge *e* has an associated positive length *l(e)*. Consider the evolution of a discrete character on *T* based on a stationary continuous-time Markov process from an unknown root state *X_ρ_* to the leaf-states (by stationarity, the prior distribution of *X_ρ_* is *π*).

We will assume that this model follows the ‘equal input model’ on *k* >1 states, with equilibrium vector *π*. In the case where *k* = 2, this class includes any stationary continuous-time Markov process; for *k* = 4 this model is known as the Felsenstein 1981 (F81 or Tajima-Nei) model (2, 19). An important property (28) of this model, is that it is equivalent to the model in which events (called *resamp/ing events* here) occur at a constant rate *γ* along the edges of the tree, and when such an event occurs the state at that point is replaced by a state chosen from the equilibrium distribution *π* independently of the original state (thus the state may or may not actually change). Conditional on the vector *π*, the transition rate *r* (i.e. the rate at which states change to different states) is related to the rate of these resampling events according to the identity:

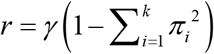

We regard *π* as an unknown quantity to be estimated (i.e. as a random variable having some distribution). The transition rate *r* is also a random variable and we will let *μ* be the expected rate of transition. Thus 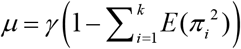 and so *μ* and *γ* are proportional to each other. Consider the following information measures as a function of *μ*. Let *X_L_* denotes the states observed at the leaves of *T*, and let:

- *I_ρ_*(*μ*) = *I* (*X_ρ_; X_L_*) be the mutual information between the state at the root vertex *ρ* and the states observed at the leaves of *T*;
- *I_π_*(*μ*) = *I* (*π; X_L_*) be the mutual information between the equilibrium distribution (*π*) for the model and the states observed at the leaves of *T*.

We can now state our first theorem (details and full proof in SI).

#### Theorem 1

For any phylogenetic tree with any number of leaves, I_ρ_(μ) is a monotone decreasing function of μ (with limit 0 as μ → ∞). By contrast, I_π_(μ) is bounded above by a monotone increasing function φ(μ), which agrees with I_π_(μ) at μ = 0 and as μ → ∞. The latter limit corresponds to the highest possible information that can be obtained with n samples drawn from π, corresponding to a multinomial distribution (given by a tree with very long edges).

The proof of both parts of Theorem 1 applies the classical Data Processing Inequality (DPI) from information theory (29), but in different ways. Recall that the DPI states that if *X, Y* and *Z* are any three random variables (not necessarily real-valued), and if *X → Y → Z* is a Markov chain, then *I(X; Z)* is less or equal to both *I(X; Y)* and *I(Y; Z)*. Moreover, unless the associated process *X → Z → Y* is also a Markov chain, then these inequalities are strict. To show that *I_ρ_(μ)* is a monotone decreasing function of *μ* we establish a more general result (allowing each edge to have its own expected transition rate) and then examine the impact of increasing this expected rate on any given edge. A (probabilistic) coupling argument, together with an application of the DPI, leads to the claimed monotonicity. For the second part of Theorem 1 (concerning *I_π_(μ))*, we consider the more informative (but unobservable) process *Q* in which one knows all the resampling events and the transitions within the tree (not just the states at the leaves). Let *φ(μ)=I(π,Q)* be the mutual information between *π* and this more informative process. Using the DPI we show that *I_π_(μ) <φ(μ)*. A further application of the DPI to the more informative process *Q* shows that *φ(μ)* is a monotone increasing function of *μ*, and the claims about the values of *φ(μ)* at *μ* = 0 and as *μ→ ∞* then follow.

### Statement and sketch proof of Theorem 2

Notice that the estimation error curves in the simulations (Fig. 2) appear to come down as *n* increases. However, it is not at all clear whether they would continue to decrease towards zero or would instead converge to some non-zero value. We show that Yule trees with fixed heights allow for asymptotically precise estimation of both the root state and the relative rates as the number of leaves become large. To simplify the calculations, we increase the speciation rate *λ* (as this grows, the number *n* of leaves is a random variable that tends to infinity). In Theorem 2 we allow more general models than the equal input model (as assumed in Theorem 1), encompassing most models used in molecular phylogenetics, including the HKY model used in our simulations.

We state Theorem 2 as follows (details and full proof in SI).

#### Theorem 2

For any continuous-time evolutionary model with positive rate matrix R, the ancestral root state, and the rate matrix R (i.e. both the relative rates and the global rate μ) can both be estimated with an error converging to zero on a Yule tree, as the number of leaves tends to infinity. However, this is not possible for other tree shapes such as the star and Kingman coalescent trees.

To show that the root state can be accurately estimated, the primitive method of maximum parsimony (MP) (2–3) suffices (even though it is less accurate than maximum likelihood). The proof that MP is consistent here combines two ideas: first we apply a (probabilistic) coupling argument which shows that it is enough to establish the result for an associated 2-state process; we then investigate this simpler process by deriving and analyzing a system of non-linear differential equations (analogous to ref. 17).

To show that the entries in the rate matrix *R* can also be consistently estimated we consider an estimation method based on 3-leaf pendant subtrees. While such a method is not likely to be optimal (e.g. maximum likelihood surely performs better) it is nevertheless sufficient to establish the theorem, and its simplicity allows for a tractable mathematical analysis that would be difficult for more complicated methods. We deal with 3-leaf pendant subtrees rather than just 2-leaf pendant subtrees (‘cherries’, commonly used to estimate models from sequence data) for two reasons. First, it allows us to consider more general Markovian processes (in particular, we need not assume the Markovian process is time-reversible). Second, even for time-reversible models, an approach based on cherries only works if the leaves are very far from the root (so that the frequencies of states is at (or very close to) equilibrium); in our setting, the tree has fixed height, and so generally the distribution of states amongst the leaves will not be very close to the equilibrium distribution (as observed in the simulations, a major difference with sequences where the equilibrium distribution is well approximated by the state frequencies among the sites, due to stationarity).

For each pair of not necessarily distinct states *i* and *j*, we say that a 3-leaf pendant subtree *(ab)c* is of type *ij* if leaf *c* and one of the remaining leaves (*a* or *b*) have state *i* and the other leaf (from the pair *a, b*) has state *j*. We will also say that *(ab)c* is of type *i* if it is of type *ik* for some *k* (including the case *k* = i), and that *(ab)c* is *typical* if its height is no more than twice its expected height (in a Yule tree). For each pair of distinct states *i, j*, let *N_ij_* denote the number of typical 3- leaf trees of type *ij* and *N_i_* denote the number of typical 3-leaf pendant subtrees of type *i*.

Define *L_i_* to be twice the sum of the heights of the cherries of the typical 3-leaf pendant subtrees of type *i*. Our proof uses the following simple estimator of the transition rates in the rate matrix *R*: for any two (distinct) states *i, j*, let 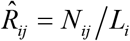. We show that 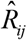 is a statistically consistent estimator of *R_ij_* as *λ* grows. The proof uses the distribution of the number and height (30–31) of 3-leaf subtrees in a Yule tree, together with further asymptotic arguments to show that 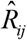 converges in probability to *R_ij_*.

For the second part of Theorem 2, first suppose that *T_n_* is a star tree. Then neither the ancestral root state, nor the equilibrium vector *π* can be estimated accurately, even as *n→∞*. More precisely, the following non-identifiability result holds. One can switch the root state to any other one other state, and adjust the parameter *π* and resampling rate to give an identical probability distribution on the data that such a tree generates regardless of how large *n* is (details are provided in the SI). This is a strongly negative result, even if biological trees are not purely star trees. With star-like trees corresponding to rapid radiations, estimations are likely to be very difficult.

Next suppose that *T_n_* is a tree generated by the Kingman coalescent. For any value of *μ* > 0, the state at the root of *T_n_* has an error that does not decrease to zero as *n→∞*. The proof relies on a well-known property (24) of the coalescent tree *T_n_*. the shorter of the two edges incident with the root of *T_n_* has an exponential distribution with a mean that is asymptotic (as *n→∞*) to *l*/2, where *l* is the height of the tree. A simple coupling argument (32) then shows that with probability at least *p* > 0 (where *p* is independent of *n* but dependent on *μ, l* and the rate matrix *R*) the states at the leaves are independent of the root state of *T_n_*, and so the error in inferring the root state does not tend to zero as *n* grows.

### Simulation protocol

To explore the behaviour of evolutionary models that are more complex and realistic than F81 and the equal input models used in Theorem 1, we performed computer simulations using the HKY model (2, 25). We generated Yule and coalescent trees with a number *n* of tips equal to 100 and 1,000. These trees were rescaled to have a total height of 1.0. Then, we simulated the evolution of a 4-state character according HKY with *κ* (transition/ transversion ratio) equal to 4.0, and *π_i_* equilibrium frequencies equal to 0.15, 0.35, 0.35, and 0.15, for A, C, G, and T, respectively. The HKY rate matrix was normalized as usual (i.e. the expected number of changes along a branch of length 1.0 was set to 1.0) and then multiplied by the global rate *μ* with values equal to 1/16, 1/8, 1/4, 1/2, 1, 2, 4, 8, and 16. For each of the tree models (Yule, coalescent) and *μ* values, 1,000 trees and data sets were generated with *n* = 100, and 500 with *n* = 1,000 for computing time reasons. For each data set we jointly estimated using the maximum likelihood principle (ML) the *μ* and *κ* parameters, the four *π_i_* equilibrium frequencies, and the ancestral character state at the tree root. The latter was inferred using the MAP (maximum a-posteriori) principle (i.e. the predicted state corresponded to the maximum of the posteriors among the four states), which is known to be optimal (26). The estimation procedure was performed in three steps:

1. As the HKY parameters were unknown, we first used the Jukes and Cantor (JC) model (2) to obtain a rough 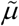 estimate of *μ* by ML. Then, the tree was rescaled by multiplying all branch lengths by 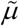.
2. The resulting rescaled tree was given to PhyML (33) along with the tips values, to estimate the *κ* and *π* parameters (corresponding to KappaML and FreqML curves in Fig. 2). It has been demonstrated in a number of studies (e.g. (34)) that the estimation of evolutionary model parameters remains accurate with approximate trees, as we have here regarding the branch lengths that are rescaled using 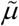(instead of *μ* that is unknown).
3. These ML-estimates of *κ* and *π* were used to jointly infer the root state (RootMLfull in Fig. 2) and obtain a better ML-estimate of *μ* assuming an HKY model (GlobalRateML in Fig. 2). To quantify the loss of accuracy induced by the approximate estimation of the model parameters (*μ*, Kand *π*), we also estimated the root state with the model parameter values used to generate the data (RootMLtrue in Fig. 2).

A difficulty with ML-based estimations of *μ*, is that when the mutation rate is low, the root and tip states tend to be identical, and then *μ* is estimated to be zero. Similarly, with high rate *μ* is often estimated to be infinite. In both cases the estimation of the other parameters and root state becomes impossible (at least using a standard ML implementation, as PhyML). Thus, we imposed the constraint: 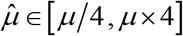, where *μ* is the true value and 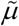 the estimate. This constraint was used with both JC-based (Step 1, 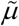) and HKY-based (Step 3) estimations of *μ*.

To quantify the estimation error, for the numerical estimates (*μ, κ* and *π*) we measured the average over all data sets of the relative absolute error (e.g. with 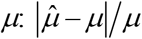, and the error of the four frequencies was further averaged over the four states. For the root state we simply measured the frequency of the prediction errors. For comparison with the more accurate ML approach, a rough estimate of the equilibrium frequencies was also obtained by counting the number of state occurrences at the tree tips (FreqTips in Fig. 2). The error of this quick estimator (used in many ML software programs, but with sequences, not a single character) is clearly higher than FreqML, corresponding to the fact that with low and moderate *μ* the tip state frequencies do not reach the equilibrium probabilities. We also compared the estimations of the *π_i_* frequencies and root state, with those obtained with a multinomial with *n* trials drawn using the same nucleotide probabilities as in tree-based simulations. This multinomial model is equivalent to the tree model when *μ* is very large and/or the pending branches are extremely long. In this condition, the tips state values do not bring any information on the root state (*I_ρ_*= 0), while the *I_π_* information on *π* is as high as possible with *n* tips/trials (see Theorem 1). The root prediction error (RootMultin in Fig. 2) is then nearly equal to 0.65 (MAP returns C and G states with ~0.5 probability each, and both have a prediction error of 0.65). The frequency estimation error was computed by simulations (FreqMultin in Fig. 2, ~0.148 and ~0.047 with *n* = 100 and 1,000, respectively). As expected RootMultin ≈ RootML ≈ 0.65 with = 16, and FreqMultin ≈ FreqML ≈ FreqTips with *μ* = 16 and Yule trees. Coalescent trees have much shorter pending branches, and the convergence of FreqML and FreqTips toward FreqMultin is slower.

All software programs -but PhyML- used to perform the simulation study were implemented in Common Lisp and are available on request. We used the version 3.3.20170530 of PhyML available from https://github.com/stephaneguindon/phyml.

## Supporting information

Supplementary Information - Mathematical proofs

## Acknowledgements

Thanks to Vincent Lefort for his help in running PhyML with large-scale simulated data. This work was supported by the INCEPTION project (PIA/ANR-16-CONV-0005, OG).

