## Supplementary Information - Mathematical proofs for "A Darwinian Uncertainty Principle"

### Mathematical Results

Some parts of the *Methods* section are repeated here for ease of readability. Reference numbering is as listed in the main manuscript.

#### 1. Statement and proof of Theorem 1

Let $T$ be a rooted phylogenetic tree (not necessarily binary), for which each edge $e$ has an associated positive length $l(e)$. Consider the evolution of a discrete character on $T$ based on a stationary continuous-time Markov process from an unknown root state $X_\rho$ to the leaf-states (by stationarity, the prior distribution of $X_\rho$ is $\pi$). We will assume that this model follows the 'equal input model' on $k > 1$ states, with equilibrium vector $\pi$. In the case where $k = 2$, this class includes any stationary continuous-time Markov process; for $k = 4$, this model is known as the Felsenstein 1981 (F81 or Tajima-Nei) model (2, 28).

An important property of this model, is that it is equivalent to the model in which events (called *resampling events* here) occur at a constant rate $\gamma$ along the edges of the tree, and when such an event occurs the state at that point is replaced by a state chosen from the equilibrium distribution $\pi$ independently of the original state (thus the state may or may not actually change) (28, 32). Conditional on the vector $\pi$, the transition rate $r$ (i.e. the rate at which states change to different states) is related to the rate of these resampling events according to the identity $r = \gamma(1 - \sum_{i=1}^{k} \pi_i^2)$. We will sometimes regard $\pi$ as an unknown quantity to be estimated (i.e. as a random variable having some distribution). The transition rate $r$ is also a random variable and we will let $\mu$ be the expected rate of transition. Thus

$$\mu = \gamma(1 - \sum_{i=1}^{k} \mathbb{E}[\pi_i^2]),$$

and so $\mu$ and $\gamma$ are proportional to each other.

Consider the following information measures as a function of $\mu$. Let $X_L$ denotes the states observed at the leaves of $T$, and let:

- $I_\rho(\mu) = I(X_\rho; X_L)$ be the mutual information between the state at the root vertex $\rho$ and the states observed at the leaves of $T$;
- $I_\pi(\mu) = I(\pi; X_L)$ be the mutual information between the equilibrium distribution ($\pi$) for the model and the states observed at the leaves of $T$.

**Theorem 1.**

*(i) For each fixed $\pi$, the mutual information $I_\rho(\mu)$ is a monotone decreasing function of $\mu$, with $I_\rho(0) = H(\pi) = -\sum_i \pi_i \ln(\pi_i)$ (the entropy of the distribution $\pi$) and $\lim_{\mu\to\infty} I_\rho(\mu) = 0$.*

*(ii) The mutual information $I_\pi(\mu)$ is bounded above by $\varphi(\mu)$, where $\varphi$ is a monotone increasing function, with $I_\pi(0) = \varphi(0)$ and $\lim_{\mu\to\infty}(I_\pi(\mu) - \varphi(\mu)) = 0$. These coin-*

*ciding values of $\varphi(\mu)$ and $I_\pi(\mu)$ at $\mu = 0$ and as $\mu \to \infty$ are the mutual information between the random vector $\pi$ and $N$ independent samples of a multinomial distribution with the probability vector $\pi$, where $N = 1$ when $\mu = 0$ and $N = n$ as $\mu \to \infty$.*

*Proof.* The proofs of both Part (i) and Part (ii) apply the classical Data Processing Inequality (DPI) from information theory, but in different ways. Recall that the DPI states that if $X, Y$ and $Z$ are any three random variables (not necessarily real-valued), and if

$$X \longrightarrow Y \longrightarrow Z$$

is a Markov chain, then $I(X;Z)$ is less or equal to both $I(X;Y)$ and $I(Y;Z)$. Moreover, unless the associated process $X \longrightarrow Z \longrightarrow Y$ is also a Markov chain, then these inequalities are strict (29). All the chains we consider in the proof are of this type and thus we have strict inequalities.

For Part (i), it is convenient to establish a more general result by allowing each edge $e$ of $T$ to have its own resampling rate, denoted $\gamma(e)$. Thus we will let $m : E(T) \to \mathbb{R}^{>0}$ be the function that assigns the edge $e$ a corresponding resampling rate $\gamma(e)$ and we let $I_\rho(m)$ denote the corresponding mutual information between $X_\rho$ and the states at the leaves of $T$.

Now suppose that the resampling rate on a fixed edge $e$ of $T$ is increased from $\gamma(e)$ to $\gamma(e) + \delta$, where $\delta > 0$, with all the transition rates on the other edges being fixed. Let $m'$ be the corresponding assignment of rates to $T$ (so $m'$ is identical to $m$ except for edge $e$, where it adds $\delta$ to the transition rate). For the pair $(T, m)$, let $X_1$ denote the vector of observed states at the leaves of $T$ that are descended from $e$ and let $X_2$ denote the vector of observed states for the remaining leaves of $T$. Similarly, for the pair $(T, m')$, let $X_1'$ denote the vector of observed states for each leaf $T$ descended from $e$, and let $X_2'$ denote the vector of observed states for the remaining leaves of $T$. Note that increasing the transition rate on edge $e$ by $\delta$ is stochastically equivalent to leaving the transition rate unchanged and lengthening the edge $e$ by adding in an additional $\delta l(e)$ length (this is illustrated in Supplementary Figure 1).

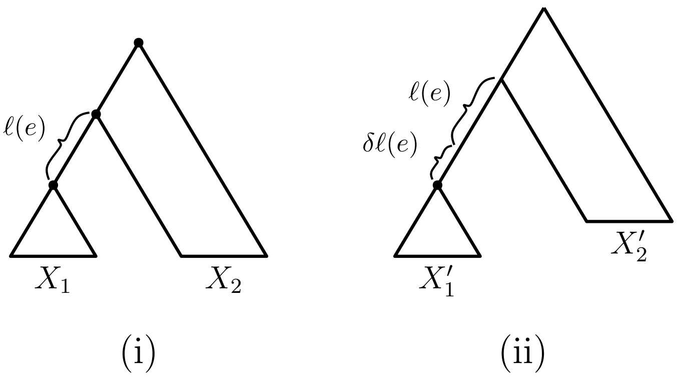


**Supplementary Figure 1**: Treating an increase in the transition rate on an edge as an increase in the edge length in the proof of Theorem 1.

Note that the probability of a resampling event in the additional segment of the edge of length $\delta l(e)$ is $1 - \exp(\delta l(e))$ Thus, if we consider the following chain of three variables:

$$X_\rho \longrightarrow (X_1, X_2) \longrightarrow (X_1', X_2'),$$

then $X_2' = X_2$ and

$$X_1' = \begin{cases} X_1, & \text{with probability } \exp(-\delta l(e)); \\ W_1, & \text{with probability } 1 - \exp(-\delta l(e)), \end{cases}$$

where $W_1$ is independent of $X_\rho$ (since a resampling event gives a state that is independent of earlier states, including that at the root).

It follows that the three-variable chain above is a Markov chain (i.e. $(X_1, X_2)$ renders $X_\rho$ and $(X_1', X_2')$ conditionally independent). Thus we can apply the DPI to obtain:

$$I(X_\rho; (X_1, X_2)) > I(X_\rho; (X_1', X_2')).$$

Now, $I(X_\rho; (X_1, X_2)) = I_\rho(m)$ and $I(X_\rho; (X_1', X_2')) = I_\rho(m')$, and so $I_\rho(m) > I_\rho(m')$. The monotonicity claim in Part (i) now follows by induction (we can simply increase the resampling rate on all the edges of $T$ one at a time) and by also noting that $\mu$ is proportional to the resampling rate $\gamma$.

For the last part of (i), notice that when $\mu = 0$, all the leaves are in the same state as the root (with probability 1) and so $I_\rho(\mu) = I(X_\rho; X_L) = I(X_\rho; X_\rho) = H(\pi)$. On the other hand, in the limit as $\mu \to \infty$, the probability that there is a resampling event on every (pendant) edge of $T$ tends to 1, and so $X_L$ is (asymptotically) independent of $X_\rho$, which gives $\lim_{\mu\to\infty} I_\rho(\mu) = 0$.

For Part (ii), we first state a simple combinatorial lemma. Given a set $P$ consisting of points on the edges of $T$, and a point $p \in P \cup \{\rho\}$, let $S_p$ be the set of leaves $x \in X$ for which there is a path from $p$ to $x$ that does not pass through any other point of $P$. Let $\partial P$ be those points $p$ in $P \cup \{\rho\}$ for which $S_p \neq \emptyset$.

**Lemma 1.** *Let $P$ be any subset of points on the edges of $T$. Then:*

*(i) If $p'$ is an additional point on an edge of $T$ then $|\partial(P \cup \{p'\})| - |\partial P|$ equals 0 or 1;*

*(ii) $1 \leq |\partial P| \leq n$, and if $|\partial P| < n$, then there is at least one pendant edge $e$ of $T$ for which if $p'$ is any point on $e$, then $|\partial(P \cup \{p'\})| = |\partial P| + 1$.*

*Proof. Part (i):* If each directed path from $p'$ to the leaf set $X$ passes through at least one point in $\partial P$ then $\partial(P \cup \{p'\}) = \partial P$ and so these two sets have the same size. Similarly, if there is a point $p \in \partial P$ for which every directed path from $p$ to the leaf set $X$ passes through a point of $P \cup \{p'\}$ then $\partial(P \cup \{p'\}) = ((\partial P) - \{p\}) \cup \{p'\}$ and so once again $|\partial(P \cup \{p'\})| = |\partial P|$. Otherwise, $\partial(P \cup \{p'\})$ is the disjoint union of $\partial P$ and $\{p'\}$ and so $|\partial(P \cup \{p'\})| = |\partial P| + 1$.

*Part (ii):* We have $1 \leq |\partial P|$ since if $P$ is empty, then $\rho \in \partial P$; the upper bound follows because $\{S_p : p \in \partial P\}$ is a partition of $X$ and so has at most $|X|$ blocks, combined with the observation that the function $p \mapsto S_p$ is one-to-one from the set $\partial P$ to $\{S_p : p \in \partial P\}$. For the second part, if $|\partial P| < n$, there is a point $p \in \partial P$ for which $S_p$ contains at least two leaves, say $x, x'$ (recall that the sets $\{S_p : p \in \partial P\}$ form a partition of $X$). Let $p'$ be any point on the pendant edge incident with $x'$. Then $|\partial P \cup \{p'\}| = |\partial P| + 1$. $\square$

We will now take $P$ to be the set of points (on the edges of $T$) where resampling events occur. Note that $\partial P$ depends only on $\mu$ (i.e not on $\pi$). If $\mu$ is low, we expect the size of $\partial P$ to be 1, whereas with large $\mu$, it is expected to be $n$ (i.e. at least one sampling event per pendant edge).

For a point $p \in \partial P$, let $Y(p)$ be the state that is resampled, which is drawn from the vector $\pi$ (in the case where $p = \rho$, then $Y(p)$ is the state at the root, which is also drawn from $\pi$), and let $\mathcal{P}$ be the vector of pairs $[(p, Y(p)) : p \in \partial P]$. Since $\mathcal{P}$ completely determines the random variable $X_L$ (the states at the leaves of $T$), we have the following Markov chain:

$$\pi \longrightarrow \mathcal{P} \longrightarrow X_L.$$

Since $I_\pi(\mu) = I(\pi; X_L)$, the DPI gives:

$$I_\pi(\mu) \leq I(\pi; \mathcal{P}).$$

Next, let $\mathcal{Q}$ be the multi-set of states $[Y(p) : p \in \partial P]$. Notice that the entries in $\mathcal{Q}$

consist of $|\mathcal{Q}|$ samples of states drawn i.i.d. according to the probability distribution $\pi$. Moreover,

$$I(\pi; \mathcal{P}) = I(\pi; \mathcal{Q}),$$

because the location of the points in $P$ and therefore in $\partial P$ is independent of the random variable $\pi$. We will take

$$\varphi(\mu) = I(\pi; \mathcal{Q}), \tag{1}$$

as the function described in the statement of Part (ii). It remains to show that $\varphi(\mu)$ is strictly monotone increasing with $\mu$, and with a limit as described in the theorem. First, let $N \in \{1, 2, \ldots, n\}$ be the random variable $|\partial P|$ (which is also the size of the multi-set $\mathcal{Q}$). We now adopt a similar strategy to the proof of Part (i). That is, we allow each edge $e$ to have its own resampling rate $\gamma(e)$ and we consider effect of increasing this value on an arbitrary edge $e$ from $\gamma(e)$ to $\gamma(e) + \delta$ for $\delta > 0$, and let $N'$ be the size of the resulting set $\partial P$. By Part (i) of Lemma 1, we have:

$$N' = N + B,$$

where the random variable $B$ is either 0 or 1, and the event $B = 1$ has a strictly positive probability that does not depend on $\pi$. We now apply a simple coupling argument. Let $\mathcal{Z} = (Z_1, Z_2, \ldots, Z_N)$ be a sequence of $N$ sample of states drawn i.i.d. according to the probability distribution $\pi$, and let $\mathcal{Z}' = (Z_1, Z_2, \ldots, Z_N, Z_{N+1})$, where $Z_{N+1}$ is a further independent state sampled according to $\pi$. Then either $\mathcal{Z} = \mathcal{Z}'$ (when $B = 0$ occurs) or else $\mathcal{Z}$ is $\mathcal{Z}'$ with the last sample excluded (when $B = 1$ occurs). Thus $\mathcal{Z}$ can be obtained from $\mathcal{Z}'$ by deleting the last entry, with a certain positive probability, or leaving it intact; the probabilities of these two outcomes do not depend on $\pi$.

Thus we can apply the DPI to the Markov chain $\pi \longrightarrow \mathcal{Z}' \longrightarrow \mathcal{Z}$ to deduce that $I(\pi; \mathcal{Z}') > I(\pi; \mathcal{Z})$. Moreover, since any ordering of a multi-set of samples is independent of $\pi$, we have:

$$I(\pi; \mathcal{Q}) = I(\pi; \mathcal{Z}), \text{ and } I(\pi; \mathcal{Q}') = I(\pi; \mathcal{Z}'),$$

where $\mathcal{Q}'$ is the multi-set of states on $T$, with $e$ having an elevated resampling rate. Thus $I(\pi; \mathcal{Q}') > I(\pi; \mathcal{Q})$, and therefore if we increase the $\gamma(e)$ values across all the edges of $T$ one at a time, and recall that $\mu$ is proportional to $\gamma$ (and invoke Eqn. (1)), we see that if $\mu < \mu'$, then $\varphi(\mu) < \varphi(\mu')$.

Regarding the last two claims in Part (ii), notice that as $\mu \to \infty$, $|\partial P|$ converges in probability to $n$ (by Part (ii) of Lemma 1), since there will be a resampling event on every pendant edge of $T$ with a probability converging to 1. This corresponds to $n$ i.i.d. samples from a multinomial distribution with probability vector $\pi$ for the limiting values of both $I_\pi(\mu)$ and $\varphi(\mu)$ as $\mu \to \infty$. At the other extreme ($\mu = 0$), we have $P = \emptyset$ (and thus $\partial P = \{\rho\}$), so all the leaves of $T$ are in the same state as the root (with probability 1) corresponding to 1 sample from this multinomial distribution for $I_\pi(0)$ and $\varphi(0)$.

□

### 2. Statement and proof of Theorem 2

For Theorem 2, we can generalise the model and allow any stationary, reversible, continuous-time Markov process (involving any number of states) for which the rate matrix $R$ has strictly positive off-diagonal entries (i.e. $R_{ij} > 0$ for all $i, j$ with $i \neq j$). This model includes the equal input models studied in the previous section (and Theorem 1), as well as most of the models used in molecular phylogenetics, including the HKY model used in our simulations.

The estimation error curves in the simulations (Fig. 2 of the main manuscript) appear to come down as $n$ increases, but it is not at all clear whether they would continue to decrease towards zero or would instead converge to some non-zero value.

We show that Yule trees with fixed heights allow for asymptotically precise estimation of both the root state and the relative rates as the number of leaves become large. To simplify the calculations we increase the speciation rate $\lambda$ (as this grows, the number $n$ of leaves is a random variable that tends to infinity). We first show that the root state can be accurately estimated; indeed for this purpose the primitive method of maximum parsimony (MP) (2,3) suffices (even though it is less accurate than maximum likelihood).

**Theorem 2.**

*(i) Consider a Yule tree $\mathcal{T}$ grown for time 1 at speciation rate $\lambda$, and $\mu$ be the overall transition rate. For any value of $\mu \geq 0$, the probability that MP correctly infers the true root state of $\mathcal{T}$ from the leaf states of this tree is bounded below by a function $f_\mu(\lambda)$ (independent of $R$), which converges to 1 as $\lambda \to \infty$.*

*(ii) For any $\mu > 0$, each of the entries in the (actual, non-normalized) rate matrix $R$ can be estimated with arbitrarily high precision as $\lambda \to \infty$.*

*(iii) Suppose that $T_n$ is a star tree with $n$ leaves and known height. Consider the equal input model for any root state value $i$, strictly positive equilibrium vector $\pi$ and resampling rate $\gamma > 0$. Then for any other root state value $j \neq i$ there is a different equilibrium vector $\pi' \neq \pi$ and a resampling rate $\gamma'$ that produces the same distribution on data at the leaves of $T_n$ for all $n$. Hence neither the root state nor equilibrium vector $\pi$ can be inferred accurately in the limit as $n \to \infty$.*

*(iv) For (Kingman) coalescent trees the accuracy with which the root state can be estimated does not converge to zero as $n \to \infty$.*

*Proof of Theorem 2(i):*

We first require a combinatorial lemma. Essentially, this lemma says that if we change all but one of the states (say, state $i$) in a multi-state character to a new 'dummy' state (denoted $\neg i$) to obtain a binary character, then if $i$ is a most parsimonious root state for this binary character on a tree, it is also most parsimonious state for the original multi-state character on that tree.

**Lemma 2.** *Let $T$ be a rooted binary phylogenetic $X$–tree with root vertex $\rho$, let $f : X \to S$ be an evolved character with arbitrary discrete state space $S$, and select any state $i \in S$, and let $\neg i$ denote a new state different from all the states in $S$.*

*Suppose that $\tilde{f} : X \to \{i, \neg i\}$ satisfies the property that $\tilde{f}(x) = i \Rightarrow f(x) = i$. Then:*

*(i) If $i$ is a most parsimonious state at $\rho$ for $\tilde{f}$, then $i$ is a most parsimonious state at $\rho$ for $f$.*

*(ii) If $i$ is the unique most parsimonious state at $\rho$ for $\tilde{f}$, then $i$ is the unique most parsimonious state at $\rho$ for $f$.*

*Proof.* Let $\mathrm{FS}(v)$ be the Fitch set (in the first pass of the Fitch algorithm (19) of the vertex $v$ for the character $f$ on $T$ and let $\tilde{\mathrm{FS}}(v)$ be the corresponding Fitch set for the character $\tilde{f}$ on $T$. We wish to show that that (i) $i \in \tilde{\mathrm{FS}}(v) \Rightarrow i \in \mathrm{FS}(v)$ and (ii) $\tilde{\mathrm{FS}}(v) = \{i\} \Rightarrow \mathrm{FS}(v) = \{i\}$.

We prove both these statements simultaneously by induction on $n = |X|$, the number of leaves of $T$. Properties (i) and (ii) clearly hold when $n \leq 2$ so suppose they both hold for all rooted binary phylogenetic trees with $n \leq k$ leaves ($k \geq 2$), and that $T$ is a tree with $k + 1$ leaves.

Consider the two maximal proper subtrees $T_1, T_2$ of $T$ obtained by deleting the root vertex $\rho$ and its incident edges. Let $\mathrm{FS}_1, \mathrm{FS}_2$ be the Fitch sets of the roots of $T_1$ and $T_2$

for the restriction of $f$ to each subtree and let $\tilde{\mathrm{FS}}_1$ and $\tilde{\mathrm{FS}}_2$ be the corresponding Fitch sets when $f$ is replaced by $\tilde{f}$. Now, suppose that $i \in \tilde{\mathrm{FS}}(v)$. Since $\tilde{f}$ is binary, this implies that either (a) $\tilde{\mathrm{FS}}_1 = \tilde{\mathrm{FS}}_2 = \{i, \neg i\}$ or (b) that $\tilde{\mathrm{FS}}_j = \{i\}$ for some $j \in \{1, 2\}$. If we now apply the induction hypothesis, Case (a) gives $i \in \mathrm{FS}_1$ and $i \in \mathrm{FS}_2$ which implies that $i \in \mathrm{FS}(v)$, whereas in Case (b) we have $\mathrm{FS}_j = \{i\}$ which again implies that $i \in \mathrm{FS}(v)$. This establishes that property (i) holds for $T$.

Now suppose that $\tilde{\mathrm{FS}}(v) = \{i\}$. Since $\tilde{f}$ is binary, this implies that either (c) $\tilde{\mathrm{FS}}_1 = \tilde{\mathrm{FS}}_2 = \{i\}$ or (d) $\tilde{\mathrm{FS}}_k = \{i\}$ and $\tilde{\mathrm{FS}}_l = \{i, \neg i\}$ for $\{k, l\} = \{1, 2\}$. Again applying the induction hypothesis, Case (c) gives $\mathrm{FS}_1 = \mathrm{FS}_2 = \{i\}$, which implies that $\mathrm{FS}(v) = \{i\}$, whereas in Case (d) we have $\mathrm{FS}_j = \{i\}$ and $i \in \mathrm{FS}_k$, which again implies that $\mathrm{FS}(v) = \{i\}$. This establishes property (ii) for $T$.

In summary, the assumption that properties (i) and (ii) hold for all values of $n$ up to $k$ implies that both properties hold for $n = k+1$. This completes the proof of the induction step, and thereby the lemma. □

Returning to the proof of Part (i) of Theorem 2, we use a coupling argument. We also start the Yule tree with a single lineage (rather than starting with two lineages) - this allows a simpler analysis, and since we show that the starting state can be estimated with accuracy tending to 1 as $\lambda \to \infty$ then the state at the first split can be estimated with even greater accuracy (though the difference converges to zero, since the length of this first edge tends to zero as $\lambda$ grows).

Let $\mu_i$ be the transition rate out of state $i$, and let $\nu = \max_i\{\mu_i\}$. Then $\mu = c\nu$ for some constant $c \in [0, 1]$. Now let $\alpha$ denote the state at the vertex $\rho$ (of out-degree 1), and consider the associated Markov process on $\mathcal{T}$ on the state space $\{\alpha, \beta\}$ in which $\alpha$ changes to $\beta$ at the rate $\nu$, and where $\beta$ is an absorbing state (i.e. it has zero rate of reverting to state $\alpha$). By Lemma 2, the probability that MP correctly infers the true root state from the leaf states under the original model is at least as high as it is for this derived 2-state process; moreover this holds regardless of the state $\alpha$ at the root (since $\mu_\alpha \leq \nu$ for all states $\alpha$).

Now let $S(t), D(t)$ and $E(t)$ be the probabilities that the Fitch set is $\{\alpha\}$, $\{\beta\}$ and $\{\alpha, \beta\}$ respectively, under the 2-state process described, on the tree $\mathcal{T}_t$ (where $0 \leq t \leq 1$) evolved for time $t$ from an initial lineage (the Fitch set of a vertex that has out-degree 1 is the Fitch set of its descendant vertex, which is either a leaf or a vertex of out-degree 2).

Note that $S(t) + D(t) + E(t) = 1$. An infinitesimal argument (extending the start of the initial lineage by $\delta$) similar to that for the (quite different) *symmetric* 2-state model (in Theorem 2.3 of (17)) shows that:

$$\begin{aligned}
S(t+\delta) &= \lambda\delta(S(t)^2 + 2S(t)E(t)) + (1 - \lambda\delta - \nu\delta)S(t) + o(\delta),\\
D(t+\delta) &= \lambda\delta(D(t)^2 + 2D(t)E(t)) + (1 - \lambda\delta - \nu\delta)D(t) + \nu\delta + o(\delta),\\
E(t+\delta) &= \lambda\delta(E(t)^2 + 2S(t)D(t)) + (1 - \lambda\delta - \nu\delta)E(t) + o(\delta).
\end{aligned}$$

For example, the first term on the right of the equation for $S(t+\delta)$ allows for a speciation event to occur in the first $\delta$ period of time (on the initial single lineage), while the second term allows for no speciation and no transition event to occur in the first $\delta$ period of time (notice that a transition event contributes to the equation for $D(t+\delta)$). The $o(\delta)$ terms consider the possibility of more than one event in the first $\delta$ period of time.

This leads to the following (quadratic) system of differential equations, where $\zeta = \nu/\lambda$

and $S = S(t), D = D(t), E = E(t)$:

$$\begin{aligned}\frac{dS}{dt} + (1+\zeta)S &= S^2 + 2SE,\\ \frac{dD}{dt} + (1+\zeta)D &= D^2 + 2DE + \zeta,\\ \frac{dE}{dt} + (1+\zeta)E &= E^2 + 2SD,\end{aligned}$$

subject to the initial initial condition for this system is $S(0) = 1, D(0) = E(0) = 0$.

We can further simplify this to a system of two equations in two variables ($S$ and $E$) by replacing $D$ in the third equation by $1 - S - E$. Then, by similar techniques to those employed in (17) (based on dynamical systems arguments, and the solution of the system of quadratic equations for the stationary system) the monotone decreasing function $S(t)$ has a non-zero limit at $t \to \infty$ of

$$s = 1 - \frac{1}{3}[1 + 2\zeta - \sqrt{1 - 14\zeta + \zeta^2}],$$

provided that

$$\zeta < 7 - \sqrt{48} \approx 0.0718.$$

Since $S(t) \geq s$ for $t = 1$, and applying Lemma 2 as discussed above, we can take as our choice of the function $f_\mu(\lambda)$ in Theorem 2, $f_\mu(\lambda) = 1 - \frac{1}{3}[1 + 2\zeta - \sqrt{1 - 14\zeta + \zeta^2}]$ for $\zeta = \nu/\lambda = \mu/c\lambda$ (recall from above that $\mu = c\nu$) and since $f_\mu(\lambda) \to 1$ as $\lambda \to \infty$ we establish Part (ii) of Theorem 2.

Supplementary Figure 2 shows the behaviour of $S(t)$ (determined by numerical solution of the differential equation system above, using MAPLE) for values of $\zeta$ below and just above the critical value.

□

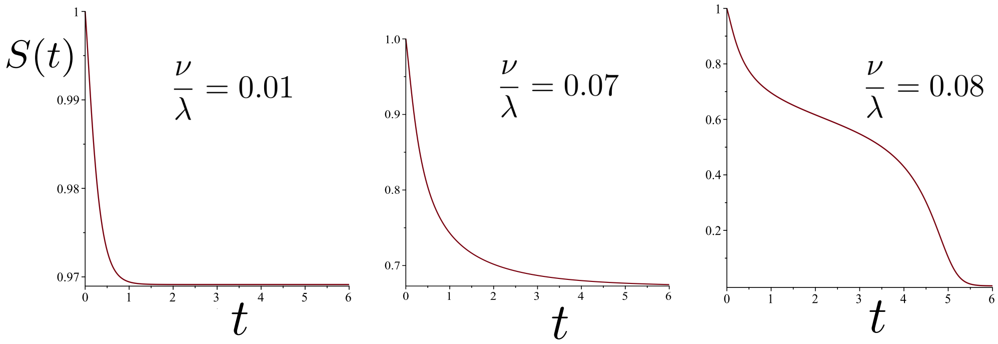


**Supplementary Figure 2**: The behaviour of $S(t)$ for values of $\zeta = \nu/\lambda$ below and just above the critical value $7 - \sqrt{48} \approx 0.0718$, showing the transition to a loss of information concerning the root state. Note that the vertical axis of these graphs all start at different heights (so $s$ is close to 0.97 when $\mu/\lambda = 0.01$).

*Proof for Theorem 2(ii):*

Our proof considers an estimation method based on 3-leaf pendant subtrees. While such a method is not likely to be optimal (e.g. maximum likelihood surely performs better) it is nevertheless sufficient to establish the theorem, and its simplicity allows for a tractable mathematical analysis that would be difficult for more complicated methods.

We deal with 3-leaf pendant subtrees rather than just 2-leaf pendant trees ('cherries') for two reasons. First, it allows us to consider more general Markovian processes (in particular, we need not assume the Markovian process is time-reversible). Second, even for time-reversible models, an approach based on cherries only works if the leaves are very far from the root (so that the frequencies of states is at (or very close to) equilibrium); in our setting, the tree has fixed height, and so generally the distribution of states amongst the leaves will not be very close to the equilibrium distribution (this is observed in the simulations).

The edges of these 3-leaf pendant subtrees, as well as the edges leading into them, can be shown (31) to be exponentially distributed with mean proportional to $\frac{1}{2\lambda}$. We will refer to a 3-leaf pendant subtree as being *typical* if its height (distance from the present as indicated by $h'$ in Fig. 3) is less than twice its expected height. Note that at least half of the 3-leaf pendant subtrees are typical, independently of $\lambda$ (since if $X$ is any non-negative random variable then Markov's inequality gives $\mathbb{P}(X \geq x) \leq \mathbb{E}[X]/x$ and setting $x = 2\mathbb{E}[X]$ gives $\mathbb{P}(X \leq 2\mathbb{E}[X]) \geq \frac{1}{2}$).

Now, for each pair of not necessarily distinct states $i$ and $j$ , we say that a 3-leaf pendant subtree with leaves $(ab)c$ is of *type* $ij$ if leaf $c$ and one of the remaining leaves ($a$ or $b$) have state $i$, and the other leaf (from the pair $a, b$) has state $j$. We will also say that $(ab)c$ is of *type* $i$ if it is of type $ik$ for some $k$ (including the case $k = i$).

Also, for each pair of distinct states $i, j$, let $N_{ij}$ denote the number of typical 3-leaf trees of type $ij$ and let $N_i$ denote the number of typical 3-leaf pendant subtrees of type $i$.

Define $L_i$ to be twice the sum of the heights of the cherries of the typical $3-$leaf pendant subtrees of type $i$ (the height of the cherry $l$ is indicated in Supplementary Figure 3). We analyse the following simple estimator of the transition rates in the rate matrix $R$. For any two (distinct) states $i, j$ let:

$$\hat{R}_{ij} := \frac{N_{ij}}{L_i}.$$

We show that $\hat{R}_{ij}$ is a statistically consistent estimator of the rate $R_{ij}$ in the limit as $\lambda$ grows.

To establish this, we first establish a number of preliminary results.

Let $N$ be the number of leaves of the Yule tree $\mathcal{T}$ growth for time $t = 1$ at speciation rate $\lambda$, and let $N_3$ be the number of 3-leaf pendant subtrees in $\mathcal{T}$. Conditional on $N = n$, the random variable $N_3$ has a mean of $n/6$ and a variance of order $n$ (ref. (30)) and so

$N_3/n$ converges in probability to 1/6 as $n \to \infty$. Now $N$ has a geometric distribution with mean $e^{\lambda h} = e^{\lambda}$ (recall that $h = 1$) (21). Thus,

$$\mathbb{P}(N > e^{\lambda/2}) = (1 - e^{-\lambda})^{e^{\lambda/2}} \to 1,$$

as $\lambda \to \infty$, and thus

$$\lim_{\lambda \to \infty} \mathbb{P}\left(N_3 \geq \frac{1}{7} e^{\lambda/2}\right) = 1.$$

Let $N_3'$ denote the number of typical 3-leaf pendant subtrees. Since at least half the 3-leaf pendant subtrees are typical, the last limit equation gives:

$$\lim_{\lambda \to \infty} \mathbb{P}\left(N_3' \geq \frac{1}{14} e^{\lambda/2}\right) = 1. \tag{2}$$

Next, let $\hat{f}_i$ denote the frequency of state $i$ amongst any antichain of vertices (not chosen with regard to the character) that are present between time $\frac{1}{2} \leq t \leq 1$ in $\mathcal{T}$, and let $f_i = \mathbb{E}[\hat{f}_i]$. Note that $f_i$ is simply the probability of observing state $i$ after time $t$ in a single lineage that starts at the root state (note that root state may be $i$ or some other state), in particular, $f_i$ is independent of $\lambda$, as it depends solely on the root state, the matrix $R$ and $t$. By the assumption that all rates are strictly positive, it follows that, for some value $\epsilon > 0$, all states $i$, and all $t$ between $\frac{1}{2}$ and 1:

$$f_i > \epsilon > 0. \tag{3}$$

Moreover, from results in (17) (namely, Proposition 3.1 (which shows that Yule trees are 'well-spread') and Lemma 3.2) the following convergence in probability holds for each state $i$ as $\lambda \to \infty$:

$$\hat{f}_i \xrightarrow{p} f_i. \tag{4}$$

Now consider a typical 3-leaf pendant tree, $(ab)c$, let $v$ be the most recent common ancestor of $a, b, c$, and let $w$ denote the interior vertex descended from $v$ (see Supplementary Figure 3). For $i \neq j$ we say that $(ab)c$ is of *invariant-type* $ij$ if $(ab)c$ is of type $ij$ and $v$ is also in state $i$. We also introduce some further definitions at this point. Let:

- $\tilde{N}_{ij}$ denote the number of typical 3-leaf pendant subtrees of invariant-type $ij$;
- $\tilde{N}_i$ be the number of typical 3-leaf pendant subtrees that have root vertex ($v$) in state $i$;
- $\tilde{L}_i$ be twice the sum of the heights of the cherries of the $\tilde{N}_i$ typical 3-leaf pendant subtrees that have root vertex ($v$) in state $i$.

**Claims:**

(i) For any value of $\epsilon > 0$, the following inequality holds with probability tending to 1 as $\lambda \to \infty$ for any pair of distinct states $i, j$: $\tilde{N}_{ij} > \frac{C}{\lambda^{1+\epsilon}} e^{\lambda/2}$ for a constant $C > 0$ (dependent only on $R$);

(ii) Also, as $\lambda \to \infty$, $\tilde{N}_{ij}/N_{ij} \xrightarrow{p} 1$;

For each state $i$ the following additional convergence in probability results hold as $\lambda \to \infty$:

(iii) $\tilde{N}_i \xrightarrow{p} \infty$ and $\tilde{L}_i \xrightarrow{p} \infty$;

(iv) $\tilde{N}_i/N_i \xrightarrow{p} 1$, and $\tilde{L}_i/L_i \to 1$.

To justify Claim (i), Eqns. (2), (3) and (4) imply that $\tilde{N}_i$ is at least some constant times $e^{\lambda/2}$ (with probability tending to 1 as $\lambda \to \infty$). Amongst these typical 3-leaf pendant subtrees, the height $h'$ of the subtree is bounded above by a term of order $\frac{1}{\lambda}$ (by the assumption that $(ab)c$ is 'typical') and with probability converging to 1 (as $\lambda \to \infty$) its cherry height $l$ is bounded below by a term of order $\frac{1}{2\lambda}$ times a term of order $\frac{1}{\lambda^{\epsilon}}$ (for any $\epsilon > 0$). To see why this last claim holds, observe that for any exponential random variable $X$ with mean $c/\lambda$ (for a constant $c$) the probability that $X$ exceeds $1/\lambda^{1+\epsilon}$ converges to 1 as $\lambda \to \infty$ (since $\mathbb{P}(X > 1/\lambda^{1+\epsilon}) = \exp(-c\lambda \cdot \lambda^{-1-\epsilon}) = \exp(-c\lambda^{-\epsilon}) \to 1$, as $\lambda \to \infty$).

Now, just one state transition is required within this 3-leaf pendant subtree to make it of invariant type $ij$, and so the probability of this event is lower bounded by a term of order $\frac{1}{\lambda^{1+\epsilon}}$.

To justify Claim (ii) select $0 < \epsilon < 1$ in Claim (i) and note the probability that a typical 3-leaf pendant subtree of type $ij$ is not of invariant-type has order $\frac{1}{\lambda^2}$ since at least two state transitions are required on the short edges (each of length of order $\frac{1}{\lambda}$) of this typical 3-leaf tree (again by the 'typical' property).

Regarding Claim (iii), the first part of Claim (iii) follows from Claim (i), since $\tilde{N}_i \geq \tilde{N}_{ij}$ for any $j \neq i$; the second part also follows from Claim (i) because the heights of the cherries of the typical 3-leaf pendant subtrees are each of height at least of order $\frac{1}{\lambda^{1+\epsilon}}$ (for any fixed $\epsilon > 0$) with probability converging to 1 as $\lambda$ grows.

To justify the first part of Claim (iv), let $N'_i$ be the number of typical 3-leaf pendant subtrees for which all three leaves and both interior vertices are in state $i$ (thus there is no state transition within this sub-tree). By definition, $N'_i < \tilde{N}_i$ and $N'_i < N_i$. Moreover, $N'_i/\tilde{N}_i$ and $N'_i/N_i$ both converge in probability to 1, since any typical 3-leaf pendant subtree counted by $N_i$ or $\tilde{N}_i$ that is not counted by $N'_i$ requires at least one state transition within that subtree, an event that has probability of order $\frac{1}{\lambda}$ (while having no state transition within the subtree has probability $1 - O(\frac{1}{\lambda})$). The second part of Claim (iv) now follows from the first part of this claim, together with the 'typical' property.

Returning to the proof, notice that $\tilde{N}_{ij}$ is a sum of $\tilde{N}_i$ independent random variables. Specifically,

$$\tilde{N}_{ij} = B_1 + B_2 + \cdots + B_{\tilde{N}_i},$$

where $B_m = 1$, if the $m$-th typical 3-leaf pendant subtree counted by $\tilde{N}_i$ is of invariant type $ij$ (otherwise $B_m = 0$). Now,

$$\mathbb{P}(B_m = 1) = 2R_{ij}l_m + O\left(\frac{1}{\lambda}\right) l_m, \quad (5)$$

where $l_m$ is the height of the cherry in the $m$-th subtree, and $O(\frac{1}{\lambda})$ is a term that allows for the possibility of two or more state transitions within this subtree. The factor 2 appearing in Eqn. (5) recognises that for a 3-leaf pendant subtree counted by $\tilde{N}_i$, a state transition from $i$ to $j$ in either one of the two edges of the cherry in this subtree leads to a pendant subtree of type $ij$. Since $\tilde{N}_i \to \infty$ as $\lambda$ grows (by Eqns. (3) and (4)), it follows that the random variable $\tilde{N}_{ij}$ (for $j \neq i$) has an asymptotic Poisson distribution with mean $R_{ij}\tilde{L}_i$. Note that the events $B_1, B_2, \ldots,$ are independent, since the state at the root of each 3-leaf pendant subtree of invariant type counted by $\tilde{N}_i$ is the same (namely state $i$).

Now, if $X$ is a Poisson random variable with mean $\eta L$ where $L$ is a random variable that tends to infinity as $\lambda \to \infty$ we have $X/L \xrightarrow{p} \eta$ (i.e. $X/L$ converges in probability to $\eta$ as $\lambda \to \infty$). Applying this to $L = \tilde{L}_i$ and $\eta = R_{ij}$, and noting from Claim (iii) above that $\tilde{L}_i \to \infty$ as $\lambda \to \infty$, we have the following convergence in probability: For all $i, j$ with $i \neq j$:

$$\frac{\tilde{N}_{ij}}{\tilde{L}_i} \xrightarrow{p} R_{ij}, \quad (6)$$

as $\lambda \to \infty$.

The term on the left of this limit is not directly provided by the observed states at the leaves (since we do not know which of the typical 3-leaf pendant subtrees of type $ij$ are of invariant type). However, this issue can be handled as follows. By Claims (i) and (iv) above, the ratios $\tilde{N}_{ij}/N_{ij}$ and $\tilde{L}_i/L_i$ both converge in probability to 1 as $\lambda \to \infty$. It now immediately follows from Eqn. (6) that:

$$\frac{N_{ij}}{L_i} \xrightarrow{p} R_{ij}, \tag{7}$$

as required to establish the statistical consistency of $\hat{R}_{ij}$ as an estimator of $R_{ij}$.

Notice that Eqn. (7), together with the assumption that all the off-diagonal entries in $R$ are non-zero, immediately provides the further conclusion that for all distinct states $i, j, j'$ we have:

$$\frac{N_{ij}}{N_{ij'}} \xrightarrow{p} \frac{R_{ij}}{R_{ij'}},$$

which shows that certain *relative* rates can be inferred consistently without knowing the actual branch lengths in the Yule tree.

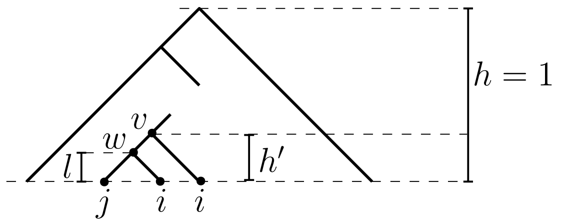


**Supplementary Figure 3**: A 3-leaf tree occurring within height $h'$ of the present, in which the outlying leaf state ($i$) also appears within the cherry of that subtree. The remaining state $j$ can be either $i$ or some other state. If $h'$ is sufficiently short, then it becomes increasingly certain that the root of the subtree will be in state $i$, and for $j \neq i$ the probability of observing $j$ at the one of the two leaves in the cherry is $2R_{ij}l$, where $l$ is the height of the cherry of the subtree.

*Proof of Theorem 2(iii):* Without loss of generality, we may assume that $T_n$ has height 1. For each state $k$, let $N_k$ be the (random variable) number of leaves in state $k$ given that the root is in state $i$. For any strictly positive equilibrium vector $\pi$ and resampling rate $\gamma > 0$, conditional on this triple of variables $(i, \pi, \gamma)$ the vector of counts $[N_k]$ has a multinomial distribution, with $n$ trials and probability vector $p = [p_k]$ where:

$$p_k = \begin{cases} \pi_i(1 - e^{-\gamma)} + e^{-\gamma}, & \text{if } k = i; \\ \pi_k(1 - e^{-\gamma}), & \text{if } k \neq i. \end{cases}$$

Let $j$ be any particular state different from $i$, and set

$$\pi'_k = \begin{cases} \frac{\pi_i(1-e^{-\gamma})+e^{-\gamma}}{1-e^{-\gamma'}}, & \text{if } k = i, \\ \frac{\pi_j(1-e^{-\gamma})-e^{-\gamma'}}{1-e^{-\gamma'}}, & \text{if } k = j, \\ \frac{\pi_k(1-e^{-\gamma)}}{1-e^{-\gamma'}}, & \text{if } k \neq i, j. \end{cases} \tag{8}$$

where $\gamma'$ is a non-zero sampling rate to be determined. It is easily checked that $\sum_k \pi'_k = 1$, and $\pi'_i, \pi'_k > 0$. Moreover, by taking $\gamma'$ sufficiently large, one can ensure that each value $\pi'_k$ (for all $k$) defined in Eqn. (8) lies in the interval $(0, 1)$. Let $p'_k$ be the probability that a leaf is in state $k$ when the triple $(i, \pi, \gamma)$ is replaced by $(j, \pi', \gamma')$. Then: $p'_i = \pi'_i(1 - e^{-\gamma'}), p'_j = \pi'_j(1 - e^{-\gamma'}) + e^{-\gamma'}$ and $p'_k = \pi'_k(1 - e^{-\gamma'})$ for all $k \neq i, j$. It can now be checked (from Eqn. (8)) that $p'_i = p_i, p'_j = p_j$ and $p'_k = p_k$ (for all $k \neq i, j$), as required.

*Proof of Theorem 2(iv):* Since we are assuming that the off-diagonal entries of $R$ are strictly positive, let $r_j = \min_{i \neq j} R_{ij} > 0$. It can be shown (32) that the process with substitution matrix $R$ (on any tree $T$) is stochastically equivalent to a process in which the following two transition processes proceed simultaneously and independently along the edges of $T$. Given the current state $i$:

(T1) transition to state $j$ with rate $r_j$;

(T2) transition to state $j$ with rate $R_{ij} - r_j$.

Consider the event $E$ that (T1) occurs on each of the two edges of $T_n$ incident with the root. Conditional on event $E$ the states at the leaves of $T_n$ are then rendered independent of the root state. Moreover, since the shorter of the two edges incident with the root of $T_n$ has an exponential distribution with a mean that has an asymptotic limit (as $n \to \infty$) of $l/2$ where $l$ is the height of the tree, it follows that event $E$ has a probability that is bounded bellow by a value $p > 0$ that depends only on $l, \mu$ and $R$ (but not $n$). Thus, with probability at least $p$ the root state is independent of the leaf states in $T_n$, regardless of how large $n$ is.
